# *Agrobacterium*-mediated Genetic Transformation and Plant Regeneration from Cotyledons in Philippine Eggplant (*Solanum melongena* L.) Acc. ‘PH 11424’

**DOI:** 10.1101/2023.07.12.548781

**Authors:** Mark Gabriel S. Sagarbarria, John Albert M. Caraan, Patrick G. Lipio, Ian Bien M. Oloc-oloc, Kazuo N. Watanabe, Desiree M. Hautea

## Abstract

Eggplant (*Solanum melongena* L.) is one of the most important vegetables grown and consumed in the Philippines; hence, continuous breeding programs are vital to maintain the supply of this economically important crop. This study demonstrates the first successful *Agrobacterium-*mediated genetic transformation and plant regeneration of a Philippine eggplant cultivar, ‘PH 11424’ also known as ‘Mistisa’. Cotyledons from two-week old seedlings were used as explants, which were transformed with disarmed *Agrobacterium tumefaciens* strain LBA4404 harboring a binary vector for CRISPR/Cas9 expression and hygromycin phosphotransferase (*HPT*), an antibiotic selection marker. Growth of shoot primordia from the agro-infected explants was observed during selective culture with 7.5 ppm hygromycin, which indicated an initial transformation success. The putatively transformed shoot primordia were then transferred to an elongation medium. The elongated and hygromycin-resistant shoots were allowed to develop roots in hormone-free medium supplemented with hygromycin, and subsequently acclimatized under greenhouse conditions. The entire process took at least 5 – 6 months. Of the total 585 agro-infected explants, the regeneration efficiency of rooted shoots was 5.1%. Successful transformation was confirmed by polymerase chain reaction (PCR) amplification of *Cas9*. Acclimatized plants tested positive for the transgene. The transgenic eggplants successfully reached maturity, flowered, and set seed. These results demonstrate a working *Agrobacterium-*mediated transformation and plant regeneration protocols using cotyledons as explants in a Philippine eggplant genotype. These biotechnology tools are critical for the successful application of genetic engineering (GE) and new breeding techniques (NBTs) in eggplant crop improvement.

## INTRODUCTION

Eggplant (*Solanum melongena* L.) is one of the most important and popular vegetable crops grown and consumed in the Philippines. It is a high-value vegetable crop, with its value reaching 6-billion pesos in 2020 (Philippine Statistics Authority (PSA) 2021). It is widely consumed and prevalent in the Filipino diet. Biotechnological applications such as genetic engineering (GE) and new breeding techniques (NBTs) enable the development of improved eggplant varieties, which promises great economic benefit. To hasten the efficiency and process of plant breeding, transgenic approaches have been employed since the mid-1990s to specifically introduce foreign DNA into plant genomes, producing crops that confer traits such as insect resistance and herbicide tolerance (Hartung and Schiemann 2014). Crops harboring the insecticidal protein Cry, *Bt* corn and *Bt* eggplant have been developed, with the latter recently approved for commercial propagation in the Philippines last October 18, 2022. While the *Bt* eggplant technology was developed through GE, conventional crossing was performed to produce the Philippine *Bt* eggplant, which were crosses between the original event produced in India by the Maharashtra Hybrid Seed Company (Mahyco) and Philippine cultivated varieties (Shelton 2021). Plant transformation and regeneration are critical steps in the application of GE and NBTs for crop improvement. However, until now there is no transformation system reported for Philippine eggplant genotypes.

*Agrobacterium*-mediated transformation in eggplant has been reported numerous times (Rotino and Gleddie 1990; Arpaia et al. 1997; Eck and Snyder 2006; Pratap et al. 2011; Subramanyam et al. 2013; Yesmin et al. 2014). However, most investigations used genotypes from India or Europe. This is important as genotype is one of the most important factors for determining the capacity of plant tissue to form somatic embryos and then to regenerate into a whole plant (Sidhu et al. 2014). In this paper we report a simple and efficient procedure for the *Agrobacterium*-mediated transformation and regeneration of a Philippine eggplant cultivar ‘PH 11424’ with *A. tumefaciens* harboring a binary vector that confers a Cas9 expression cassette and resistance to the antibiotic hygromycin.

## MATERIALS AND METHODS

### Time and Place of the Study

This study was carried out at the Institute of Plant Breeding (IPB) Biosafety Level 2 (BL2) laboratory and greenhouse at the University of the Philippines Los Baños from February 2022 to April 2023.

### Permissions

All experiments were conducted in accordance with the Joint Department Circular No. 1 Series of 2016 for contained use. The Department of Science and Technology Biotechnology Committee (DOST-BC) issued the corresponding Biosafety Permit No. 2020-0324. The biosafety permit conditions were complied with throughout the conduct of every experiment and associated greenhouse and laboratory activities under the supervision of the UPLB Institutional Biosafety Committee and the DA-BPI Central Post-Entry Quarantine Station.

### Plant material and growth conditions

Eggplant seeds of Philippine variety ‘PH 11424’ also known as ‘Mistisa’ were obtained from the National Plant Genetic Resources Laboratory, Institute of Plant Breeding, UPLB. This modern-bred cultivar was chosen based on its baseline regeneration efficiency from cotyledon explants (around 30%), which was the highest among the other genotypes screened in previous regeneration experiments done without transformation. Plant materials for transformation were grown *in vitro*. Seeds were surface sterilized with a 10% commercial bleach and 0.02% Tween 20 solution for 20 minutes with continuous mixing. Seeds were rinsed thrice with sterile distilled water. Afterwards, seeds were dried on sterile filter paper and placed into sterile bottles containing half-strength Murashige and Skoog (MS) medium with 1.5% sucrose, pH 5.8, and solidified with 0.45% sucrose. The cultures were maintained in a growth room at 25°C and under 16:8 photoperiod.

### Bacterial strain and growth conditions

*Agrobacterium tumefaciens* strain LBA4404, an octopine strain with Ach5 chromosomal background (Ooms et al. 1982), carrying a modified binary vector pZH_PubiMMCas9 (Mikami et al. 2015) was used. pZH_PubiMMCas9 contains an expression cassette for Cas9 under the control of the maize ubiquitin promoter, and an expression cassette for *hygromycin phosphotransferase* (*HPT*), under control of the 35S promoter from the cauliflower mosaic virus. Bacteria were grown at 28°C in YEB medium (5 g/L beef extract, 1 g/L yeast extract, 5 g/L peptone, 5 g/L sucrose, 0.5 g/L MgCl2, pH 7.0) with shaking at 200 rpm until they reached an OD_600_ equal to 1.0 (around 20 – 24 hours). Bacteria were resuspended in ½ strength MS medium with 1.5% sucrose and 100 μM acetosyringone and was used for co-cultivation.

### Stable transformation and plant regeneration

*Agrobacterium*-mediated transformation of eggplant (*Solanum melongena* L. cv. Mistisa) cotyledons was carried out. After agro-infection, an established eggplant regeneration protocol using direct organogenesis and zeatin riboside (García-Fortea et al. 2020) was performed with slight modifications to eliminate remaining bacteria and select for transformed cells. All culture conditions were at 25°C and under 16:8 photoperiod. Cotyledons from two-week old seedlings were excised to obtain explants around 0.5 – 1 cm^2^ in size; small wounds using the scalpel tip were incised at the adaxial side. Explants were inoculated adaxial-side-down on sterile Petri dishes containing ½ strength MS medium with 1.5% sucrose, 0.45% vitro agar, 2 mg/L zeatin riboside, and 100 μM acetosyringone for two days. Afterwards, in a sterile Petri dish, explants were submerged in the *Agrobacterium* solution for 5 minutes before drying onto a sterile paper towel and transferring back to the same medium again, still adaxial-side-down, for a 3-day co-cultivation period. Explants were washed once with sterile distilled water and then once with a 300 mg/L timentin solution before drying on a sterile paper towel and transferring adaxial-side-up to selective medium (½ strength MS, 1.5% sucrose, 0.45% vitro agar, 2 mg/L zeatin riboside, 100 mg/L timentin, 7.5 mg/L hygromycin, 200 mg/L PVP40). Prior to transformation, a hygromycin sensitivity assay was done to test varying concentrations of hygromycin (0, 5, 7.5, 10 and 15 mg/L), and it was determined that 7.5 mg/L hygromycin was a sufficient concentration that did not allow regeneration of non-transformed explants in a sample size of 30 explants.

Shoot primordia were allowed to develop for 3 – 4 weeks. Explants were then transferred to elongation medium (½ strength MS, 1.5% sucrose, 0.45% vitro agar, 0.5 mg/L gibberellic acid (GA_3_), 100 mg/L timentin, 7.5 mg/L hygromycin). Shoots were allowed to develop with sub-culturing every 3 weeks to fresh elongation medium. Calli that developed were also sub-cultured; calli that already formed shoots were discarded afterwards to avoid propagating shoots from the same cell lineage. Hygromycin concentration was lowered at this stage to encourage root formation. Shoots around 2 cm in size were transferred to rooting medium (½ strength MS, 1.5% sucrose, 0.45% vitro agar, 100 mg/L timentin, 5 mg/L hygromycin). Shoots were allowed to root twice prior to acclimatization. Rooted shoots were acclimatized in ambient laboratory conditions (approx. 29°C, moderate light) for 1 week and subsequently in ambient greenhouse conditions for an additional week prior to transfer to soil. Flowers were self-pollinated by assisted pollination using forceps and bagging of flowers. Physiologically mature fruit were harvested 60 days after pollination.

### Amplification of transgenes

Approximately 200 mg of leaf discs was used to extract genomic DNA from transformed and regenerated plants (Paterson et al. 1993). The DNA pellet was resuspended in 40 – 50 μL of TE buffer and stored at -20°C until use. Transgenes were amplified via PCR using the following primers: (MMCas9-F: 5’ –GACCCGCAAGTCTGAAGAAACT – 3’; MMCas9-R: 5’-TCTTGAGGAGATCGTGGTACGT – 3’). The amplification reaction mixture consisted of 1X buffer (Invitrogen), 2mM MgCl_2_, 0.2mM dNTPs, 0.2uM of the forward and reverse primers, 1U of Taq polymerase (Invitrogen), and 10 ng of genomic DNA. The thermal profile was: 94°C for 5 min, 35 cycles of 94°C for 30 sec, 58°C for 45 seconds, and 72°C for 1 min, then 72°C for 7 min.

## RESULTS AND DISCUSSION

From a total of 585 cotyledon explants, 30 rooted shoots were regenerated resulting in a 5.1% regeneration efficiency ([number of rooted shoots / number of total explants] × 100). Direct shoot organogenesis, indicated by shoot primordia developing directly from the explants, was observed 3 to 4 weeks in selective medium (**Fig. 1A**), even after agro-infection and the presence of 7.5 mg/L hygromycin in the medium. Upon transfer to the elongation medium, shoots elongated and further developed. However, formation of white, friable calli was also observed, which proliferated in the elongation medium. Calli eventually turned brown, but some shoots were also observed to develop from these calli (**Fig. 1B**). Time for shoot development varied greatly – some shoots that developed directly from the explant tissue, elongated the earliest at 8 weeks from agro-infection. In contrast, emergence of new shoots and elongation of extant shoots derived from calli significantly took longer, at approximately 6 months. The long regeneration time from calli could be attributed to reduced organogenic potential because only gibberellic acid (GA_3_) was the hormone present in the elongation medium, and the added stress or selection pressure from the presence of hygromycin (Tran and Sanan-Mishra 2015). Furthermore, accumulated genetic and epigenetic changes characteristic in serially sub-cultured plant tissues might have contributed to the reduction of organogenesis (Gaspar et al. 1999). Shoots had formed roots in hormone-free medium supplemented with 5 mg/L hygromycin (**Fig. 1C**), and around 5 months was required to obtain fully acclimatized rooted plants. Calli-derived shoots that were additionally grown took up to 12 months to fully elongate and develop roots. After acclimatization, transformed plants were able to reach maturity, producing flowers and fruit, from which seeds were extracted (**Fig. 1D-F**).

**Figure 1.**
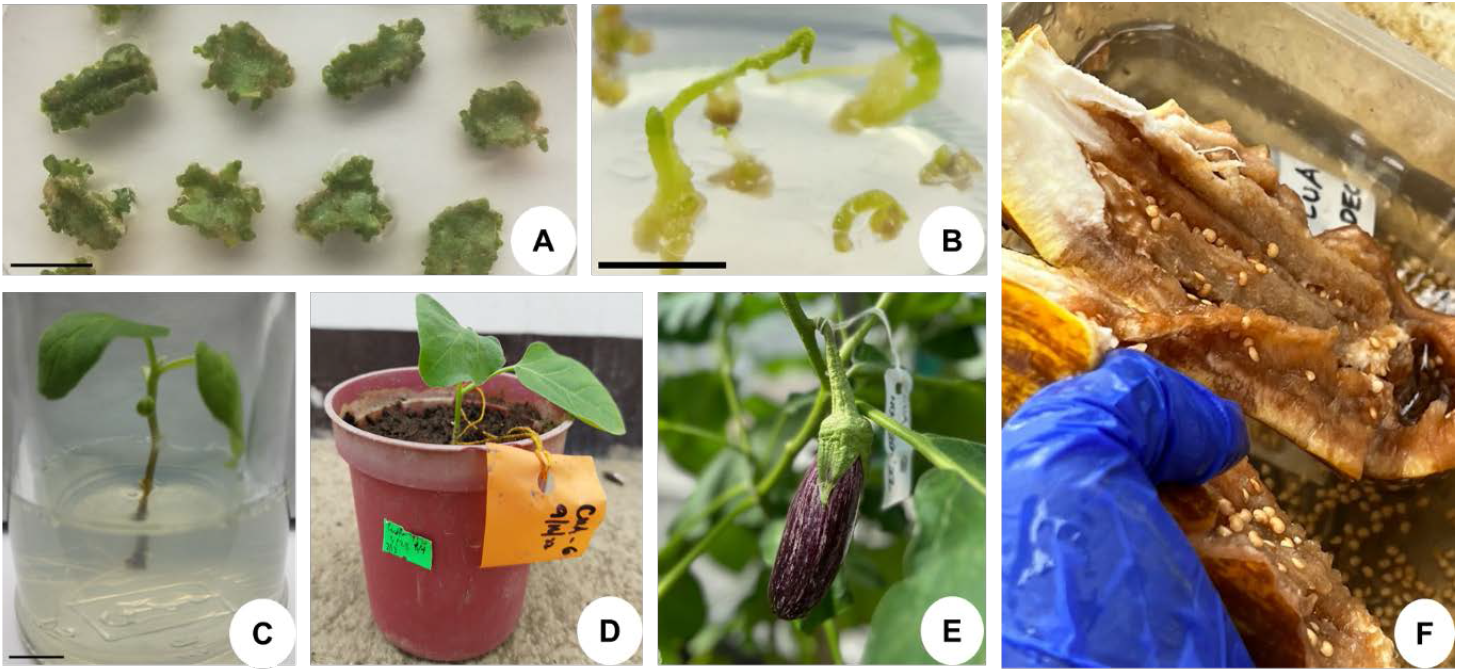
A) Shoot regeneration from cotyledon explants after *Agrobacterium* co-cultivation, in medium containing 7.5 mg/L hygromycin and 100 mg/L timentin after three weeks. Scale bar = 1 mm. B) Shoot elongation in elongation medium containing 7.5 mg/L hygromycin and 100 mg/L timentin. Scale bar = 1 mm. C) Rooting of regenerated shoots. Scale bar = 1 mm. D) Acclimatization of rooted shoots in greenhouse conditions and planting in soil. E) Fruit production of a self-pollinated flower in the transformed plants. F) Extraction of seeds from physiologically mature self-pollinated fruits.

Results of the PCR test detected positive amplification products of *Cas9* from DNA extracted from a subset of 13 acclimatized plants, while the wild-type (WT), non-transformed control of ‘PH 11424’ tested negative for the *Cas9* band (**Fig. 2**).

**Figure 2.**
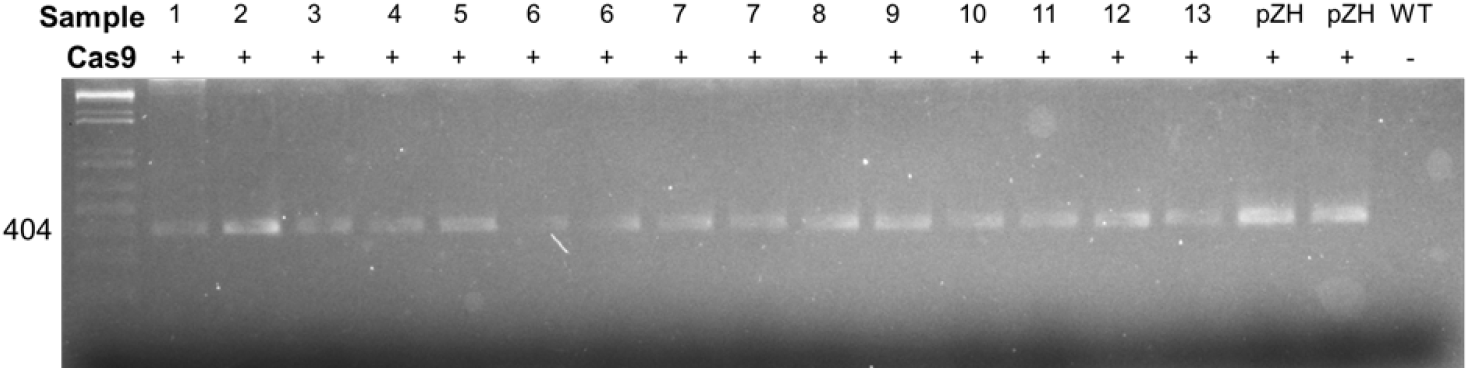
Amplification of Cas9 in a sample of 13 acclimatized plants. pZH = positive control using the binary vector pZH_PubiMMCas9 as template during PCR. WT = negative control using wild-type PH11424 DNA as template during PCR.

## CONCLUSION

These results describe a successful *Agrobacterium*-mediated transformation and regeneration protocol for a Philippine eggplant genotype. *Agrobacterium* transformed plants were able to regenerate shoots through both direct organogenesis and an intermediary callus step. The low regeneration efficiency (5.1%), which is common in many plant regeneration protocols, was otherwise offset by high transformation efficiency (i.e., consistent transgene presence) and transformant viability (i.e., reproductive capability) among acclimatized plants that developed to fully maturity. These results will provide a useful platform for future eggplant genetic engineering and genome editing studies, which can ultimately pave the way towards a more robust Philippine Agribiotechnology sector.

## ACKNOWLEDGMENT

This research was funded through the project grant to UPLB (Project Code: N91832A) by the Department of Science and Technology - Philippine Council for Agriculture, Aquatic and Natural Resources Research and Development (DOST-PCAARRD) and the matching funds provided by the Institute of Plant Breeding. The authors would like to thank the Tsukuba Plant Innovation Research Center in the University of Tsukuba (UT), Japan for providing access to the *Agrobacterium* isolates and constructs used in this study. The authors would also like to thank Ms. Rowena Frankie, Genetics Lab Technician II, for her assistance in the transformation and sub-culturing activities.

## AUTHOR CONTRIBUTIONS

MGSSagarbarria wrote the first draft of the manuscript. MGSSagarbarria, JAMCaraan, IBMOloc-oloc, and PGLipio performed tissue culture and transformation experiments. KNWatanabe and DMHautea supervised the early stages of this work. MGSSagarbarria, KNWatanabe and DMHautea contributed to study conception. All authors contributed to the revision of the manuscript and approved the final version.

## REFERENCES CITED

Arpaia S, Mennella G, Onofaro. V, Perri E, Sunseri F, Rotino GL. 1997. Production of transgenic eggplant (Solanum melongena L.) resistant to Colorado Potato Beetle (Leptinotarsa decemlineata Say). Theor Appl Genet. 95(48):329–334. doi:10.1007/s001220050567.

Eck J VAN, Snyder A. 2006. Eggplant (Solanum melongena L.). In: Wang K, editor. Agrobacterium Protocols. Vol. 343. New Jersey: Humana Press. p. 439–447.

GarcÍa-Fortea E, Lluch-Ruiz A, Pineda-Chaza BJ, GarcÍa-PÉrez A, Bracho-Gil JP, Plazas M, Gramazio P, Vilanova S, Moreno V, Prohens J. 2020. A highly efficient organogenesis protocol based on zeatin riboside for in vitro regeneration of eggplant. BMC Plant Biol. 20(1):6. doi:10.1186/s12870-019-2215-y. https://bmcplantbiol.biomedcentral.com/articles/10.1186/s12870-019-2215-y.

Gaspar T, Kevers C, Bisbis B, Franck T, Crevecoeur M, Greppin H, Dommes J. 1999. Loss of Plant Organogenic Totipotency in the Course of in vitro Neoplastic Progression. Source: In Vitro Cellular & Developmental Biology Plant. 36(3):171–181. [accessed 2023 Apr 17]. http://www.jstor.org/stable/4293333.

Hartung F, Schiemann J. 2014. Precise plant breeding using new genome editing techniques: Opportunities, safety and regulation in the EU. Plant Journal. 78(5):742–752. doi:10.1111/tpj.12413.

Mikami M, Toki S, Endo M. 2015. Parameters affecting frequency of CRISPR/Cas9 mediated targeted mutagenesis in rice. Plant Cell Rep. 34(10):1807–1815. doi:10.1007/s00299-015-1826-5.

Ooms G, Hooykaas PJJ, Van Veen RJM, Van Beelen P, Regensburg-Tuink TJG, Schilperoort RA. 1982. Octopine Ti-Plasmid Deletion Mutants of Agrobacferium tumefaciens with Emphasis on the Right Side of the T-Region.

Paterson AH, Brubabker CL, Wendel JF. 1993. Genomic DNA suitable for RFLP or PCR analysis. Plant Mol Biol Report. 11(2):122–127.

PHILIPPINE STATISTICS AUTHORITY (PSA). 2021. Major Vegetables and Rootcrops Quarterly Bulletin, January-December 2021 (Eggplant). Philippines.

Pratap D, Kumar S, Raj SK, Sharma AK. 2011. Agrobacterium-mediated transformation of eggplant (Solanum melongena L.) using cotyledon explants and coat protein gene of Cucumber mosaic virus. Indian J Biotechnol. 10(1):19–24.

Rotino GL, Gleddie S. 1990. Transformation of Eggplant (Solanum-Melongena L) Using a Binary Agrobacterium-Tumefaciens Vector. Plant Cell Rep. 9(1):26–29.

Shelton AM. 2021. Bt Eggplant: A Personal Account of Using Biotechnology to Improve the Lives of Resource-Poor Farmers. American Entomologist. 67(3):52– 59. doi:10.1093/ae/tmab036.

Sidhu MK, Dhatt AS, Sidhu GS. 2014. Plant regeneration in eggplant (Solanum melongena L.): A review. Afr J Biotechnol. 13(6):714–722. doi:10.5897/AJBX2013.13521. http://academicjournals.org/journal/AJB/article-abstract/008D21642978.

Subramanyam K, Rajesh M, Jaganath B, Vasuki A, Theboral J, Elayaraja D, Karthik S, Manickavasagam M, Ganapathi A. 2013. Assessment of factors influencing the Agrobacterium-mediated in planta seed transformation of brinjal (Solanum melongena L.). Appl Biochem Biotechnol. 171(2):450–468. doi:10.1007/s12010-013-0359-z.

Tran TN, Sanan-Mishra N. 2015. Effect of antibiotics on callus regeneration during transformation of IR 64 rice. Biotechnology Reports. 7:143–149. doi:10.1016/j.btre.2015.06.004. https://linkinghub.elsevier.com/retrieve/pii/S2215017X15000399.

Yesmin S, Sarker RH, Hoque MI, Hashem A, Islam MS. 2014. Agrobacterium Mediated Genetic Transformation of Eggplant (Solanum melongena L.) Using Cotyledon Explants. Nuclear Science and Applications. 23(1):41–46.

